# Parasitism to mutualism shift during a viral infection in Drosophila under thermal stress

**DOI:** 10.64898/2026.10.31.763142

**Authors:** Rubén González, Mauro Castelló-Sanjuán, Anamarija Butković, Hervé Blanc, Alexandros Belavilas-Trovas, Jared Nigg, Maria-Carla Saleh

**Affiliations:** Institut Pasteur, Université Paris Cité, Viruses and RNA Interference Unit, 75015 Paris, France

**Author notes:** CINBIO, Universidade de Vigo, 36310 Vigo, Spain (Rubén González).

## Abstract

Host–parasite interactions exist along a continuum from parasitism to mutualism, yet the environmental drivers governing transitions along this spectrum remain underexplored. Here we show that chronic thermal stress systematically attenuates virulence across multiple natural RNA viruses in *Drosophila melanogaster* and shifts persistent Nora virus infection into conditional mutualism, enhancing host lifespan, reproductive output, and behavioral performance without compromising viral persistence. Driven by host immunometabolic reprogramming and aging deceleration rather than viral genomic adaptation, this phenotypic switch demonstrates that thermal stress can invert the sign of biotic interactions. Our findings establish that persistent infections can expand host thermal reaction norms in animals, revealing how climate warming can fundamentally alter selection pressures and host–pathogen coevolutionary trajectories.

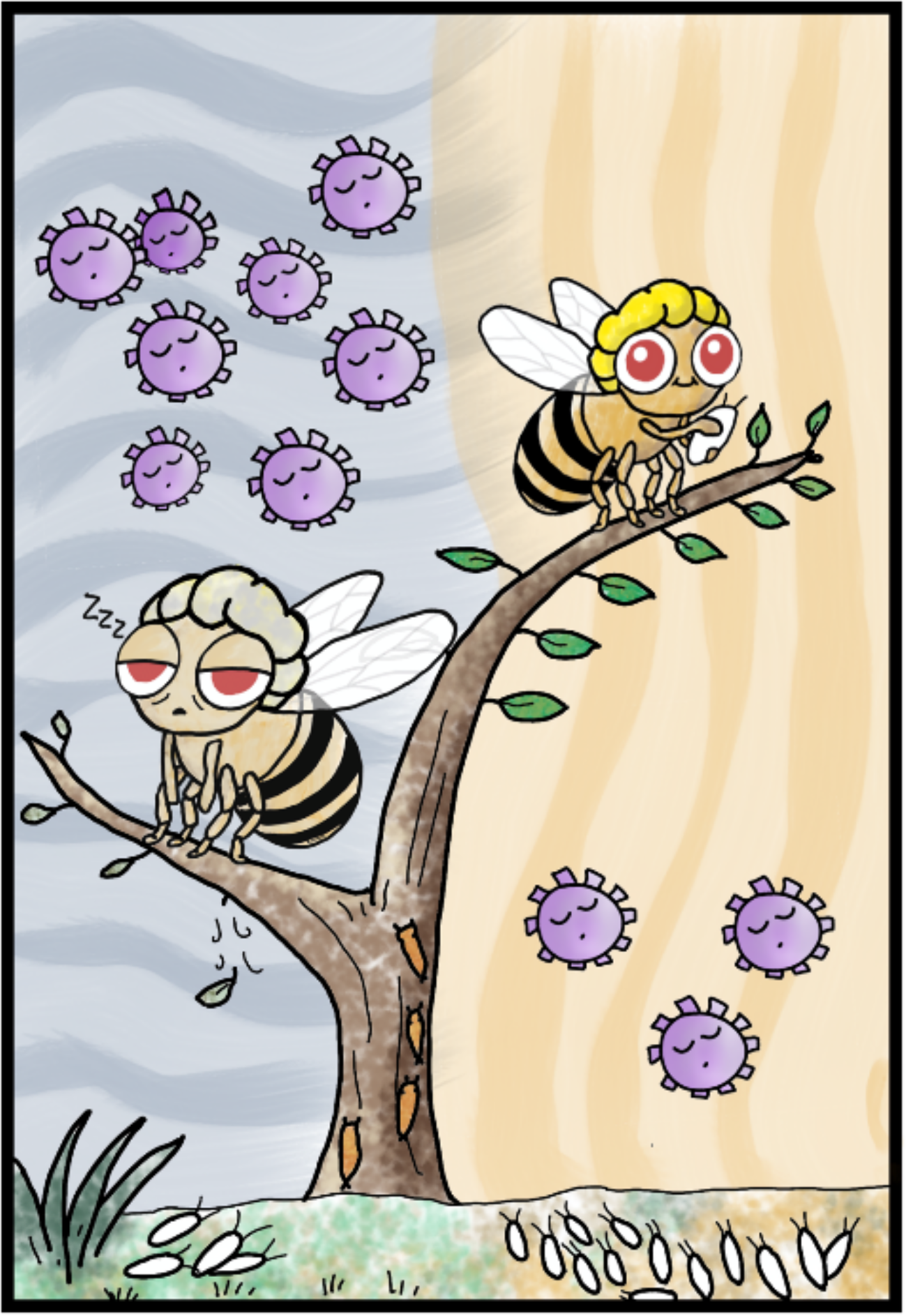

---

Host–virus interactions are traditionally viewed through the lens of pathogenesis, yet they exist along a continuum from parasitism to mutualism depending on host, viral, and environmental factors (Pagán et al., 2008; Råberg et al., 2009). In persistent viral infections, the chronic presence of the pathogen creates an extended physiological window in which environmental conditions continuously shape host fitness outcomes (Goic & Saleh, 2012; Castelló-Sanjuán et al., 2025). For ectothermic organisms, ambient temperature is the primary environmental driver governing metabolic rates, immune competence, and viral replication kinetics (Altizer et al., 2013; Hoffmann & Parsons, 1990). How thermal shifts alter the impact of infections on host fitness has direct implications for disease dynamics and host– pathogen coevolution (Thomas & Blanford, 2003; Allen & Little, 2011; Padfield et al., 2020; Duval et al., 2025). Understanding these temperature-dependent interactions is critical as ongoing climate change accelerates the frequency, intensity, and duration of terrestrial heatwaves (Perkins-Kirkpatrick & Lewis, 2020; Intergovernmental Panel On Climate Change (IPCC), 2021).

The fruit fly *Drosophila melanogaster* is a premier genetic and physiological model whose core metabolic, innate immune, and proteostatic pathways exhibit deep evolutionary conservation across metazoans (including extensive homology with mammalian systems) (Perveen 2018). This model animal allows us to dissect the fundamental principles governing host– pathogen plasticity under environmental stress. In *D. melanogaster*, persistent RNA virus infections at a standard laboratory temperature of 25°C act as classic parasites: they reduce survival, accelerate biological aging, and impair reproduction (Webster et al., 2015; Wallace & Obbard, 2024; Castelló-Sanjuán et al., 2025; González et al., 2026). Here, we tested whether an elevated temperature of 29°C, representing the upper boundary of *D. melanogaster*’s thermal niche (Hoffmann & Parsons, 1990), alters these parasitic interactions across natural positive-sense RNA viruses: Drosophila A virus (DAV), Drosophila C virus (DCV), Bloomfield virus, and Nora virus. Persistently mono-infected flies established via larval oral acquisition were maintained at 25°C or permanently shifted to 29°C following adult eclosion to systematically track fitness across the host and the virus: host survival, reproductive output, behavioral integrity, and transcriptional response, viral replication kinetics, and virus genome evolution.

To establish how chronic thermal stress reshapes host–virus fitness outcomes, we first evaluated adult lifespan under combined viral infection and elevated temperature. The 25°C survival data presented here are reproduced from Nigg et al. (2024), whereas the 29°C assays were conducted in parallel and are reported for the first time in this study. High temperature (29°C) systematically reduced baseline host lifespan, decreasing median survival (LT_50_) in uninfected flies from 63 days at 25°C to 33 days at 29°C (Fig. 1A). At 25°C, viral infections were parasitic, reducing host lifespan across all viral species with relative LT_50_ ratios ranging from 0.40 to 0.83 compared to uninfected controls (Supplementary Fig. 1). However, thermal stress at 29°C consistently attenuated the relative lifespan cost of infection across all tested viruses (Supplementary Fig. 1). To characterize this thermal modulation, we focused on Drosophila A virus (DAV, the most virulent pathogen) and Nora virus (the mildest persistent infection). In DAV-infected flies, elevated temperature reduced LT_50_ from 25 days at 25°C to 22 days at 29°C. In contrast, Nora virus infection inverted host fitness outcomes under thermal stress: while it reduced LT_50_ at 25°C (52 days vs. 63 days uninfected, relative LT_50_ = 0.82), it extended median lifespan at 29°C relative to uninfected controls (38 days vs. 33 days uninfected, relative LT_50_ = 1.15) (Fig. 1A). Thermal stress thus shifts Nora virus from a mild parasite into a conditional mutualist that buffers host lifespan against heat-induced mortality.

**Figure 1.**
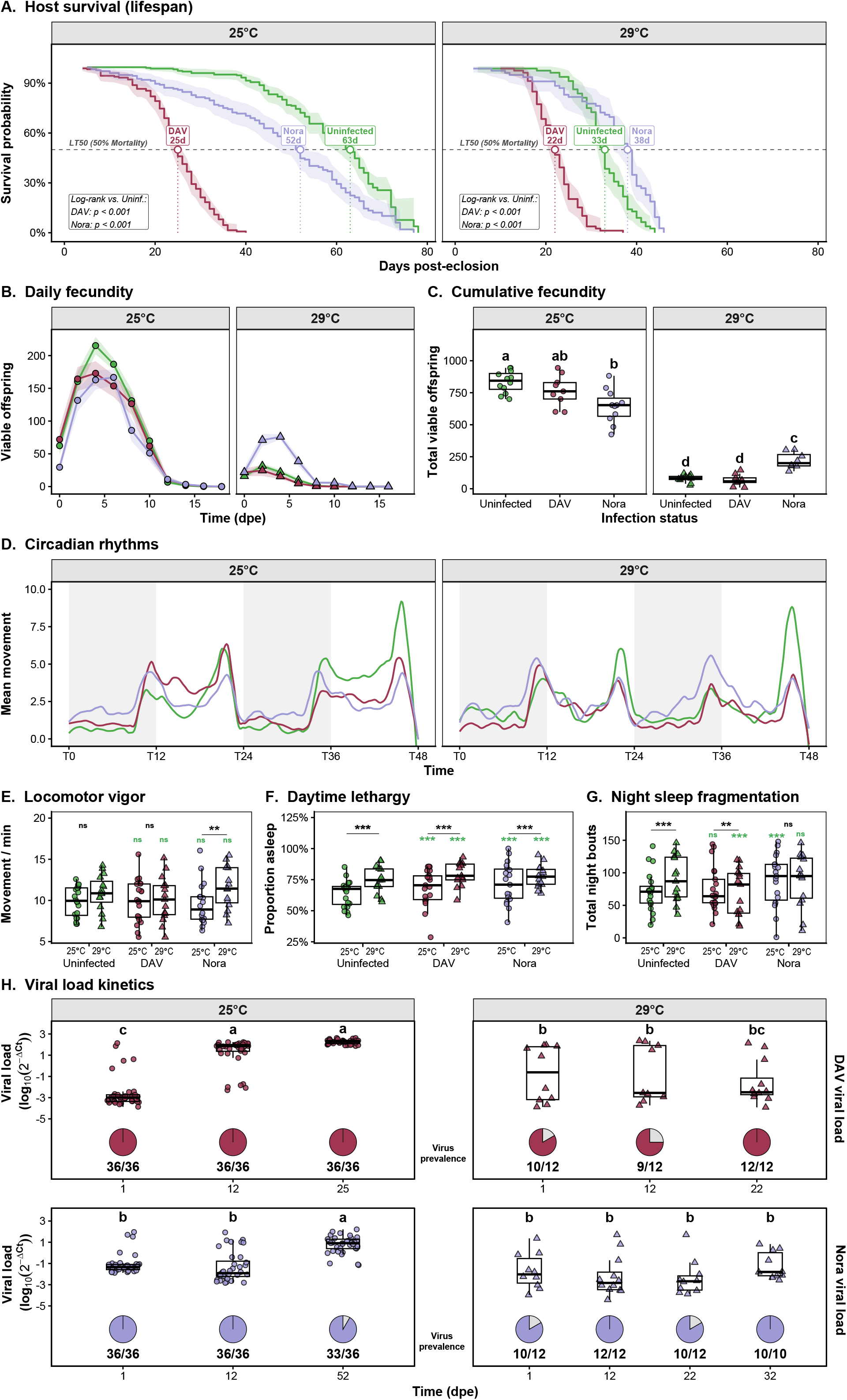
Temperature-dependent impact of persistent viral infections on host fitness, behavior, and viral load kinetics. **(A) Host survival**. Kaplan-Meier lifespan curves for uninfected controls (green), DAV-infected (red), and Nora virus-infected (purple) flies at 25°C (left panel) and 29°C (right panel). The dashed reference line marks median survival times (LT_50_), labeled in days for each treatment. Shaded regions represent 95% confidence intervals. Pairwise log-rank tests were performed to compare each viral infection against uninfected controls (*p* < 0.001). **(B) Fecundity kinetics**. Number of viable offspring (mean±SE) per population (20 females) over time (days post-adult eclosion, dpe) at 25°C and 29°C across infection conditions. **(C) Cumulative fecundity**. Total viable offspring per female across thermal conditions. Boxplots represent median and interquartile range (IQR) overlaid with individual fly population values. Statistical significance was evaluated using a linear mixed model (LMM) with experimental block as a random effect, and temperature and viral infection as interacting factors. Distinct lowercase letters indicate statistically significant differences across all temperature–treatment combinations (Tukey’s HSD post-hoc test, *p* < 0.05). **(D) Circadian rhythms**. Average locomotor activity tracks over a 48-hour continuous window (T0–T48) at 25°C and 29°C under 12:12 h light/dark cycles. **(E–G) Locomotor vigor and sleep architecture**. Quantification of locomotor vigor (movement counts per active minute; LMM) (E), daytime lethargy (proportion of time spent asleep during light hours; Binomial GLMM) (F), and night sleep fragmentation (total night sleep bout count; Poisson GLMM) (G) evaluated across the interaction of infection status and temperature, with experiment included as a random effect. Boxplots represent IQR overlaid with individual fly measurements. Green asterisks denote statistical significance compared to uninfected controls within each temperature (\**p* < 0.05, \*\**p* < 0.01, \*\*\**p* < 0.001; FDR-adjusted post-hoc tests). Horizontal brackets indicate statistically significant thermal effects within a given infection state. **(H) Viral load kinetics and prevalence**. Relative viral RNA quantification (2^-ΔCt^, normalized to *Rp49*) for DAV (top) and Nora virus (bottom) at 25°C and 29°C across time. Boxplots depict viral load distributions with individual fly measurements. Pie charts below indicate infection prevalence (infected flies / total tested). Distinct lowercase letters denote statistically significant differences across timepoints and thermal regimes (Tukey’s HSD post-hoc test, *p* < 0.05).

To evaluate whether viral mitigation of thermal stress extends beyond survival to organismal performance, we measured host fecundity and behavior under chronic viral infection at elevated temperature. While high temperature impaired overall production of viable offspring in uninfected controls (Fig. 1B, C), Nora virus infection buffered this thermal reproductive decline (Fig. 1B). This represents a phenotypic switch: although Nora virus acts as a parasite that reduces offspring production at 25°C, infected flies at 29°C produced significantly more total viable offspring than temperature-matched uninfected controls (Fig. 1C). Behavioral monitoring revealed that circadian activity architecture was preserved across all temperature and infection conditions (Fig. 1D). Baseline locomotor vigor (the mean movement counts per minute of awake flies) was unaffected by viral infection at 25°C. However, chronic exposure to 29°C increased vigor in Nora virus–infected flies (Fig. 1E). Both elevated temperature and viral infection increased daytime lethargy, with thermal stress increasing the proportion of time slept across all conditions (Fig. 1F). Finally, thermal stress significantly increased night sleep fragmentation (an indicator of impaired rest) in uninfected controls and DAV-infected flies (Fig. 1G). In contrast, Nora virus infection conferred thermal resilience against sleep disruption: although Nora virus increased baseline fragmentation at 25°C, elevated temperature induced no additional sleep fragmentation (Fig. 1G). Thus, persistent viral infection can stabilize host behavioral performance under chronic thermal stress.

To evaluate whether elevated temperature clears the infection or alters viral accumulation dynamics, we quantified viral kinetics across host lifespan. Viral titers at 29°C were restricted 50-to 100-fold compared to 25°C, with a more pronounced effect observed for DAV (Fig. 1H). This reduction in viral accumulation occurred without viral clearance, as infection prevalence remained near 100% across all time points for both viruses (Fig. 1H).

Longitudinal genomic sequencing identified no consistent directional substitutions in viral consensus sequences at 29°C, indicating that thermal restriction operates as a host-driven physiological constraint rather than requiring viral genomic adaptation (Supplementary Fig. 2). Furthermore, despite the marked reduction in DAV viral load at elevated temperatures, we detected defective viral genomes (DVGs) for DAV at 29°C in the late stages of infection. The presence of DVGs confirms that the viral RNA-dependent RNA polymerase remains actively engaged in replication rather than becoming catalytically inactive (Supplementary Fig. 3).

To establish the transcriptomic architecture underlying the observed thermal rescue, we performed longitudinal RNA sequencing of infected hosts across age and thermal regimes, evaluating differentially expressed genes (DEGs) relative to uninfected controls at the corresponding temperature. Overall, viral infection at 29°C expanded total DEG counts compared to 25°C for both DAV and Nora virus across time points (Supplementary Fig. 4A). However, cross-condition comparisons revealed a reduced DEG overlap between viruses and temperatures (Supplementary Fig. 4B). This lack of overlap suggests that the infected host response to thermal stress is mediated by pathogen-specific regulatory adjustments rather than a uniform host adaptation.

To better understand the basis of host thermal resilience, we examined the kinetic activation of heat-stress–related genes (Fig. 2A). At the standard 25°C baseline, both DAV and Nora virus infections induced a moderate upregulation of canonical chaperone genes (*Hsp70Bb, Hsp70Bc, Hsp23*) coupled with a distinct late-stage downregulation of Turandot stress-response genes (*TotA, TotC, TotM*). Under chronic thermal stress (29°C), the two viruses triggered highly divergent transcriptomic trajectories. DAV infection drove an early and sustained upregulation of *Turandot* genes alongside *GstE1*, an insect-specific oxidative stress-response enzyme, without a concurrent induction of the heat-shock protein repertoire. In contrast, Nora virus infection at 29°C suppressed the *Turandot* response throughout the different time-points but triggered a robust, late-stage upregulation of canonical heat-shock proteins (*Hsp70Ba, Hsp70Bb, Hsp70Bc, Hsp23*). These results demonstrate that DAV and Nora virus infections intrinsically modulate the host heat-stress machinery at standard temperatures. Under thermal stress, this transcriptomic rewiring diverges: Nora virus selectively upregulates the chaperone network, establishing a proteostatic profile that correlates with host thermal resilience.

**Figure 2.**
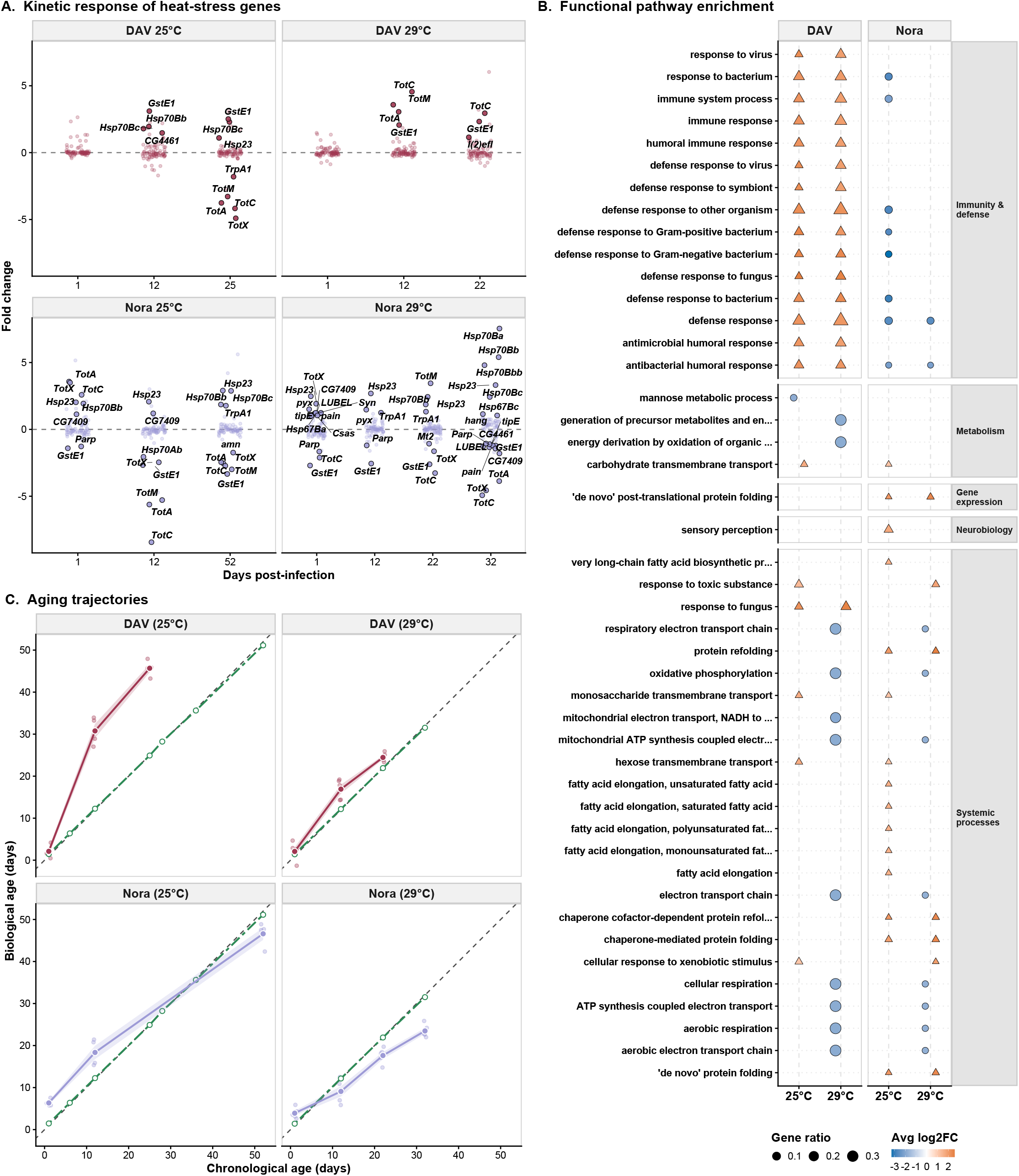
Temperature-dependent transcriptomic reprogramming, functional pathway divergence, and biological aging trajectories. **(A) Kinetic response of heat-stress genes**. Trajectories of key heat response genes (Gene Ontology GO:0009408) across days post-infection for DAV and Nora virus infections at 25°C and 29°C relative to temperature-matched uninfected controls. Faint points represent non-significant genes; highlighted bordered points denote differentially expressed genes (DEGs; *padj* < 0.05, |log_2_FC| > 1). Text labels highlight top up- and down-regulated genes at each timepoint. **(B) Functional pathway enrichment**. Gene Ontology enrichment analysis across major biological categories (immunity and defense, metabolism, gene expression, neurobiology, and systemic processes). Circle sizes represent gene ratios, and colors indicate directional expression changes (average log_2_FC; red = upregulated, blue = downregulated) for each viral infection relative to uninfected controls at the corresponding temperature (25°C or 29°C). **(C) Biological aging trajectories**. Estimated biological age (Y-axis) plotted against chronological age in days (X-axis) for DAV-infected (red) and Nora virus-infected (purple) flies compared to uninfected controls (green) at 25°C (left panels) and 29°C (right panels). Colored lines represent model trajectories; large points represent mean biological age per timepoint (n = 4 pools of 3 to 5 flies each per condition), with shaded areas denoting standard error. The dashed diagonal line indicates a 1:1 baseline correlation between chronological and biological age.

To map the functional pathways driven by this thermal transition, we conducted unbiased Gene Ontology (GO) enrichment analysis across disease states and thermal regimes (Fig. 2B). At 25°C, persistent infection by DAV or Nora virus forced sustained upregulation of energetically demanding immune defense pathways, including Toll signaling, humoral antibacterial responses, and phagocytosis. At 29°C, host immune defense was markedly attenuated during Nora virus infection. Instead, the transcriptomic profile shifted toward metabolic processes, carbohydrate and lipid transport, protein refolding, and oxidative phosphorylation (Fig. 2B).

Finally, we tested whether this persistent thermal stress response influences host biological age. At 25°C, DAV infection induced severe aging acceleration, an effect that was attenuated at 29°C. For Nora virus, infection at 25°C induced modest transcriptomic aging acceleration; at 29°C, however, Nora-infected flies exhibited a biological age significantly younger than their chronological age throughout late life (Fig. 2C). Thus, elevated temperature reverses virus-induced aging, driving a transcriptomic deceleration of biological age in Nora-infected hosts.

In summary, under chronic thermal stress, persistent RNA virus infections in *D. melanogaster* activate the host proteostatic machinery, restrict viral accumulation, and modulate host immunometabolism. Whether viral accumulation is directly restricted by chaperone hyperactivation (Merkling et al., 2015) or by temperature-induced attenuation of polymerase activity (Dalton et al., 2006) remains an open question. Regardless of the precise upstream mechanism, these physiological shifts mitigate virus-induced aging and, as observed during Nora virus infection, decelerate host aging, thereby conferring systemic resilience against environmental stress.

Our results provide direct mechanistic evidence that viral infections reshape insect thermal reaction norms and expand phenotypic plasticity under abiotic stress (González et al., 2020). Crucially, because this transition arises through host physiological plasticity rather than requiring de novo viral mutation, extreme thermal events can immediately invert the net fitness sign of infections in natural populations. While environmentally conditioned viral mutualism has been established in plants (Xu et al., 2008; González et al., 2021), our findings demonstrate that temperature-driven transitions along the parasitism–mutualism continuum represent a broad eco-evolutionary principle extending to animals. As climate warming exposes ectotherms to more frequent and prolonged heatwaves, selection may shift from host resistance toward maintaining conditional symbioses that buffer thermal stress, fundamentally altering disease dynamics, host thermal niches, and host– pathogen coevolution (Altizer et al., 2013).

## METHODS

### Fly stocks and infections

For all experiments, we used *Wolbachia*-free *w*^*1118*^ flies maintained on a standard cornmeal diet. This diet was prepared by combining 440 g inactive dry yeast, 440 g cornmeal, and 60 g agar in 6 L of osmotic water. The mixture was autoclaved and after cooling, 150 ml moldex solution (20% methylhydroxybenzoate) and 29 ml propionic acid were added as preservatives. All experiments were conducted at 25°C or 29°C under a 12:12 light:dark cycle. All fly lines were cleaned of possible persistent infections (viruses and *Wolbachia*) as described previously (Merkling & Van Rij, 2015).

Persistent viral infections were established as previously described (Nigg et al., 2022, 2024), and are the same stocks used in the work of Nigg and colleagues. Briefly, for DAV, the Australian isolate (DAV-HD, sourced from the van Rij laboratory) was injected into adult *w*^*1118*^ flies (50 nl per fly). After the injection, flies were transferred into a new clean fly food vial to contaminate the environment for three days before naïve *w*^*1118*^ flies were exposed. Persistently infected F1 progeny were isolated and maintained separately. The Nora virus-infected line originated from a *w*^*1118*^ stock persistently infected with Nora virus, provided by Dr. Stefan Ameres. Upon receipt, we repeatedly backcrossed these infected *w*^*1118*^ flies to our laboratory-maintained *w*^*1118*^ line. Confirmation that the backcrossed flies retained persistent Nora virus infection was obtained through RNA sequencing.

### Lifespan analysis

Twenty mated female flies per replicate were sorted at 1 day post-eclosion (dpe). Nine biological replicates from three independent experiments were analyzed per condition at each temperature. Males and females were co-housed for 24 hours to allow mating, then separated. Survival of females was monitored daily by counting dead flies, and flies were transferred to fresh food vials every 2 days. Fly survival was assessed from the day of adult eclosion until death. The survival curves at 25°C originate from Nigg et al. (2024), having been generated in parallel with the initial 29°C datasets presented in this work.

### Fecundity assay

Newly emerged females and males (N = 20 per sex per replicate) were co-housed for 24 hours to allow mating. Males were subsequently discarded, and mated females were transferred to fresh food vials every 2 days until egg laying ceased. Emerged adult offspring were counted up to 14 days after each laying period to ensure complete development. Cumulative lifetime fecundity was calculated by summing the normalized daily offspring per female.

### Locomotor activity and sleep architecture

Locomotor activity and sleep patterns were recorded using the DrosophilaActivity Monitoring system (DAM2, TriKinetics). Mated female flies (12 dpe) were individually loaded into glass monitoring tubes containing standard food. Flies were allowed to acclimate to the monitoring environment for 24 hours prior to data collection to eliminate handling stress. Following acclimation, activity was recorded continuously at 1-minute intervals over a 48-hour window (T0–T48) under 12:12 h light:dark cycles at 25°C or 29°C. Flies exhibiting complete immobility prior to the end of the recording window were considered dead and excluded. Sleep was defined as any continuous bout of inactivity lasting ≥5 minutes. Three behavioral metrics were quantified: (1) locomotor vigor (mean movement counts per active minute), (2) daytime lethargy (proportion of time spent asleep during light hours), and (3) night sleep fragmentation (total number of sleep bouts during dark hours).

### RNA extraction

Individual female flies were collected and homogenized using a pestle in 300 μl TRIzol reagent (Invitrogen, 15596026). RNA was extracted following manufacturer’s instructions with GlycoBlue (Invitrogen, AM9516) added during precipitation. RNA concentrations were measured using Qubit RNA BR Assay Kit (Invitrogen, Q10211).

### Viral load quantification

Four μl of RNA were treated with DNase I (Roche, 04716728001), followed by reverse transcription using random primers and Maxima H Minus Reverse Transcriptase (Thermo Scientific, EP0751). Resulting cDNA was diluted 1:10 with water. Quantitative PCR was performed using Luminaris Color HiGreen qPCR Master Mix (Thermo Scientific, K0374) on a QuantStudio 7 Flex machine (Thermo Scientific) with virus-specific primers targeting viral polymerases (Supplementary Table 1). Cycling conditions: 2 min at 50°C, 10 min at 95°C, followed by 40 cycles of 15 s at 95°C and 60 s at 60°C, with melt curve analysis. Viral loads were normalized to the housekeeping gene Rp49 and expressed as log_10_2^-ΔCt^. Samples with Ct values >35 or aberrant melt curves were considered uninfected. For pooled RNA-seq samples (5 flies per pool), mean viral loads were calculated from individual RT-qPCR measurements of the constituent flies.

### High-throughput sequencing

RNA sequencing data from 25°C was generated in Castelló-Sanjuán et al. (2025). For the 29°C samples, we performed RNA sequencing on 4 biological replicates (each replicate consisting of a pool of 5 individuals with similar viral loads, confirmed by RT-qPCR) per experimental condition. We prepared RNA-seq libraries from 15 ng of pooled RNA (3 ng per individual) using the NEBNext Ultra II RNA Library Prep Kit (New England Biolabs, E7770L) with NEBNext Multiplex Oligos (Dual Index Primers Set 1; New England Biolabs, E7600S). We performed all sequencing on an Illumina NextSeq 500 instrument using a NextSeq 500/550 High Output Kit v2.5 (75 cycles; Illumina, 20024906).

### Viral mutation analysis

Raw sequencing reads were quality-filtered and trimmed using Cutadapt to remove adapters and technical artifacts. To ensure high-quality alignments, reads shorter than 40 nucleotides were discarded, and 17 bases were strictly trimmed from the 5’ end. Trimmed reads were aligned to their respective viral reference genomes (GenBank:FJ150422.1 for DAV and GenBank:OK188768.1 for Nora virus) using HISAT2 with spliced alignments disabled. Alignments were processed, sorted, and indexed using SAMtools. High-confidence mutations were called using LoFreq, and per-position base compositions were extracted using pysamstats. To strictly isolate biological variation from sequencing noise, sub-consensus variants were filtered using bcftools to retain only those with an allele frequency (AF) ≥ 0.01. Defective viral genomes (DVGs) were identified, quantified, and classified from the sorted alignment files using DVGfinder (Olmo-Uceda et al., 2022).

### Differential expression analysis

Raw sequencing reads were mapped to *D. melanogaster* genome release dmel-r6.56 using STAR (v2.7.11b) with default parameters. Feature counting was performed using HTSeq (v0.11.2) with default settings. Differential expression analysis used DESeq2 with *padj* < 0.05 and |log_2_ fold change| > 1.0 as significance thresholds. Expression trajectories of heat-stress response genes (Gene Ontology GO:0009408) were extracted across post-infection timepoints. Functional pathway enrichment analysis of global DEGs was conducted using *clusterProfiler* (v4.10.1) with org.Dm.eg.db (v3.18.0) targeting Biological Process (BP) ontologies (*p* < 0.05, *q* < 0.05). Enriched terms were categorized into major functional domains using keyword filtering on term descriptions.

### Biological age estimation

Transcriptomic biological age was estimated using temperature-specific Elastic Net regularized regression models (*glmnet* v4.1.8). Raw count matrices for each thermal regime (25°C and 29°C) were normalized using Variance Stabilizing Transformation (VST) via DESeq2 (v1.42.1). For each temperature, an aging clock model (α = 0.5) was trained on the VST gene expression profiles of uninfected control flies across chronological age timepoints. The optimal regularization parameter (λ_min_) was selected via cross-validation using *cv*.*glmnet* (set.seed = 42). The trained model coefficients were subsequently applied to predict transcriptomic biological age for both uninfected controls and persistently infected (DAV and Nora virus) samples within their respective thermal regimes.

### Statistical analyses

All statistical modeling and data visualization were executed in R (v4.3.2) within RStudio (2026.8.0.187). Survival distributions were analyzed using Kaplan-Meier estimation and log-rank tests via *survival* (v3.8.3) and *survminer* (v0.5.0). Cumulative fecundity and locomotor vigor were evaluated using Linear Mixed-Effects Models (LMMs) via *lme4* (v1.1.37) and *lmerTest* (v3.1.3) with experimental block or monitor included as a random intercept. Daytime lethargy (proportions) and night sleep fragmentation (counts) were modeled using Generalized Linear Mixed Models (GLMMs) with Binomial (logit link) and Poisson (log link) error distributions, respectively (lme4 v1.1.37). Estimated marginal means and post-hoc pairwise comparisons were calculated using *emmeans* (v1.11.0) with Tukey’s HSD or Benjamini-Hochberg FDR corrections for multiple testing. Compact Letter Displays (CLD) were generated using *multcomp* (v1.4.28). Differential gene expression analysis was performed using *DESeq2* (v1.42.1). Functional pathway enrichment analysis was conducted using *clusterProfiler* (v4.10.1) with the org.Dm.eg.db genome annotation database (v3.18.0). Transcriptomic biological age prediction was implemented via Elastic Net regularized regression using *glmnet* (v4.1.8). Data wrangling, temporal processing, and multipanel figure assembly were performed using *tidyverse* (v2.0.0), lubridate (v1.9.4), broom (v1.0.8), *ggrepel* (v0.9.6), and *patchwork* (v1.3.0), with graphics rendered using *svglite* (v2.1.3).

## ACKNOWLEDGMENTS

We thank all members of the Saleh laboratory for helpful discussions and constructive feedback on this work. We thank Craig Cameron for advice on viral polymerases activity. We acknowledge the use of Gemini (Google) for assistance with script refinement, manuscript editing, and proofreading.

## AUTHOR CONTRIBUTIONS

Conceptualization: R.G. and M-C.S.

Methodology: R.G.

Investigation: R.G., M.C-S., A.B., J.N., H.B., A.B-T.

Visualization: R.G.

Supervision: M-C.S.

Writing—original draft: R.G.

Writing—review and editing: R.G., M.C-S., A.B.,

M-C.S.

Funding acquisition: M-C.S.

Data curation: R.G.

Project administration: R.G. and M-C.S.

Formal analysis: R.G.

## DATA AVAILABILITY

RNA sequencing data generated in this study are deposited in the NCBI Sequence Read Archive under BioProject accessions PRJNA1235228 (25°C data from Castelló-Sanjuán et al., 2025) and PRJNA1480044 (for 29°C data). All other data supporting the findings of this study are available within the paper and its Extended Data. Source data for all figures are provided as Supplementary Data files. Custom R scripts for data analysis and figure generation are available from the corresponding author upon reasonable request.

## SUPPLEMENTARY FIGURES

**Supplementary Figure 1.**
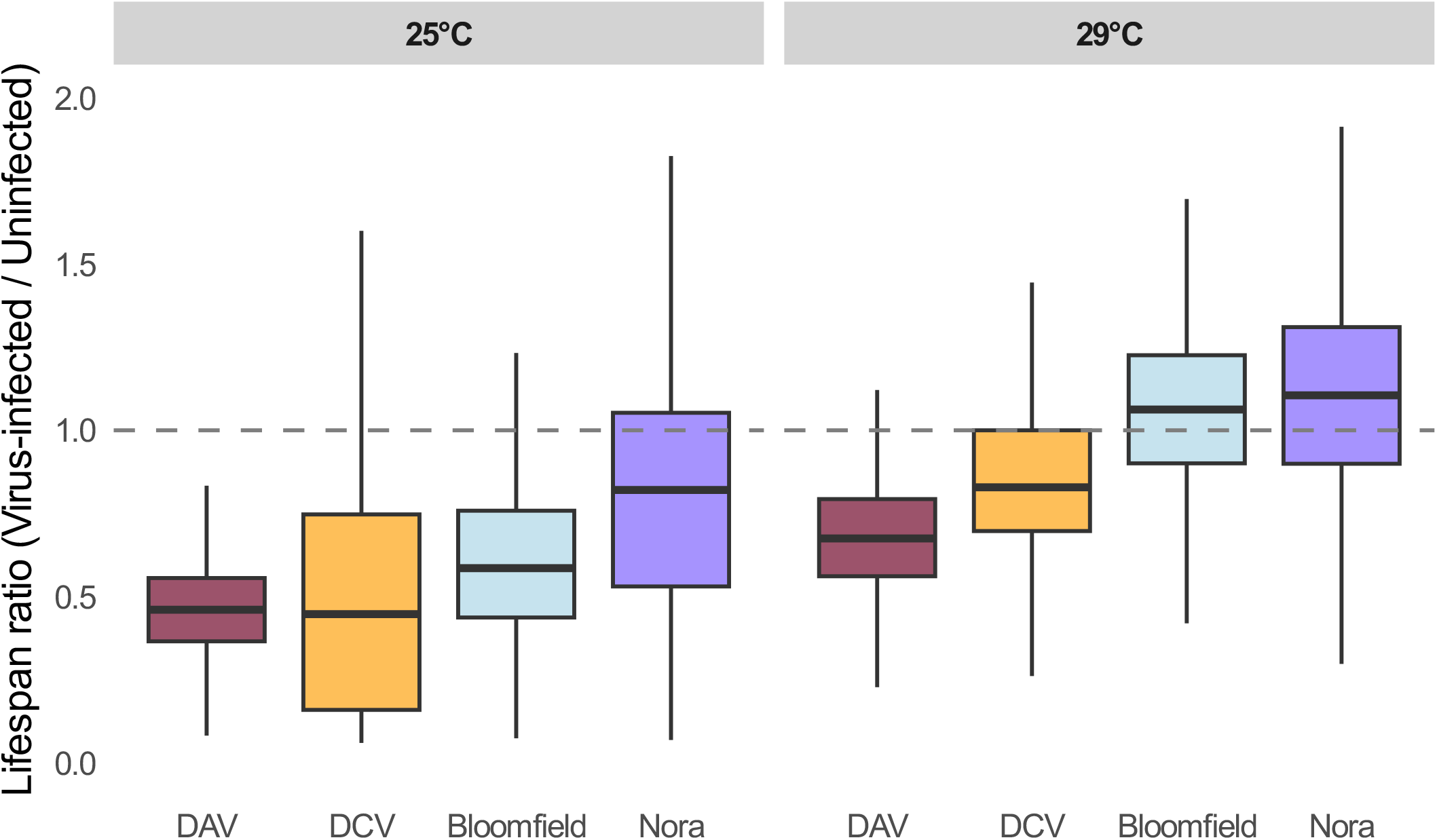
Relative host lifespan during persistent viral infections under standard and thermal stress conditions. Boxplots depict the lifespan ratio of *D. melanogaster* persistently infected with Drosophila A virus (DAV), Nora virus, Drosophila C virus (DCV), or Bloomfield virus relative to uninfected controls at 25°C and 29°C. The horizontal dashed line (y = 1) represents a lifespan equivalent to temperature-matched uninfected flies. Values below 1 indicate a relative lifespan reduction, whereas values above 1 indicate relative lifespan extension. Ratio distributions were generated using a computational resampling approach. For each condition, individual lifespans of infected females were randomly sampled and divided by the lifespan of randomly sampled uninfected females at the corresponding temperature. This procedure was iterated 1000 times per virus-temperature combination to calculate the lifespan ratios. Boxplots display the median and interquartile range of the simulated ratios, with outliers omitted for visual clarity.

**Supplementary Figure 2.**
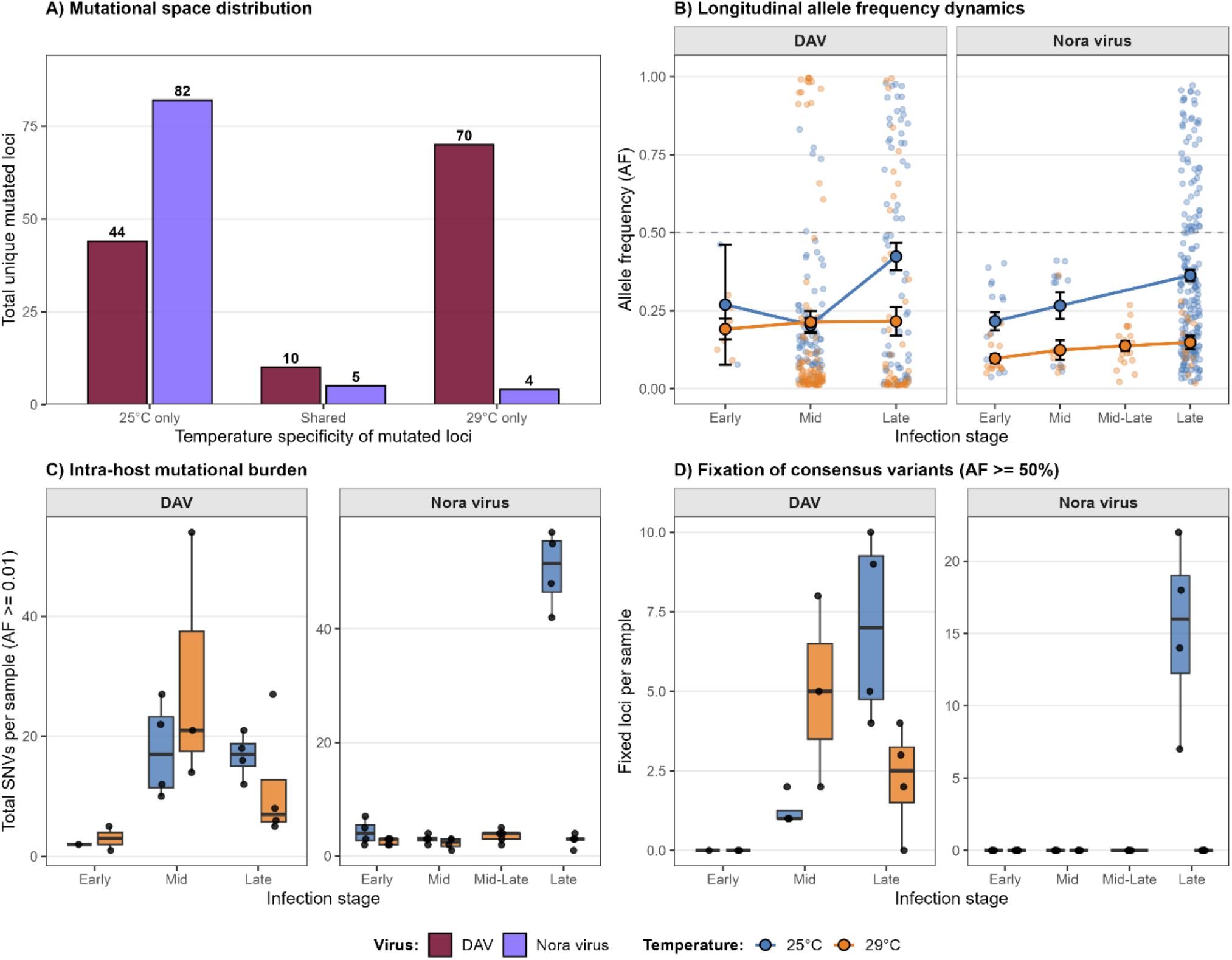
Thermal restriction limits viral accumulation without driving directional genomic adaptation in persistent virus infections. **(A)** Mutational space distribution showing the total number of unique mutated genomic loci (allele frequency, AF ≥ 0.01) classified by temperature specificity (25°C only, shared between temperatures, and 29°C only) for DAV and Nora virus. **(B)** Longitudinal allele frequency dynamics across infection stages (Early: 1 dpi; Mid: 12 dpi; Mid-Late: 22 dpi for Nora virus at 29°C; Late: 22/25 dpi for DAV at 25°C/29°C, 32/52 dpi for Nora virus at 25°C/29°C) in persistently infected flies maintained at 25°C (blue) and 29°C (orange). Points and error bars represent the mean ± standard error (SE) across mutated loci; smaller translucent points depict individual locus AF values per sample. The horizontal dashed line marks the consensus threshold (AF = 0.50). **(C)** Intra-host mutational burden quantified as the total number of single nucleotide variants (AF ≥ 0.01) per biological sample (pool of 5 flies) across infection stages. **(D)** Fixation of consensus variants quantified as the number of mutated loci with AF ≥ 0.50 per sample across infection stages, including replicates with zero fixation events. Boxplots in (C) and (D) display the median and interquartile range overlaid with black points representing each sample (pools of 5 individuals).

**Supplementary Figure 3.**
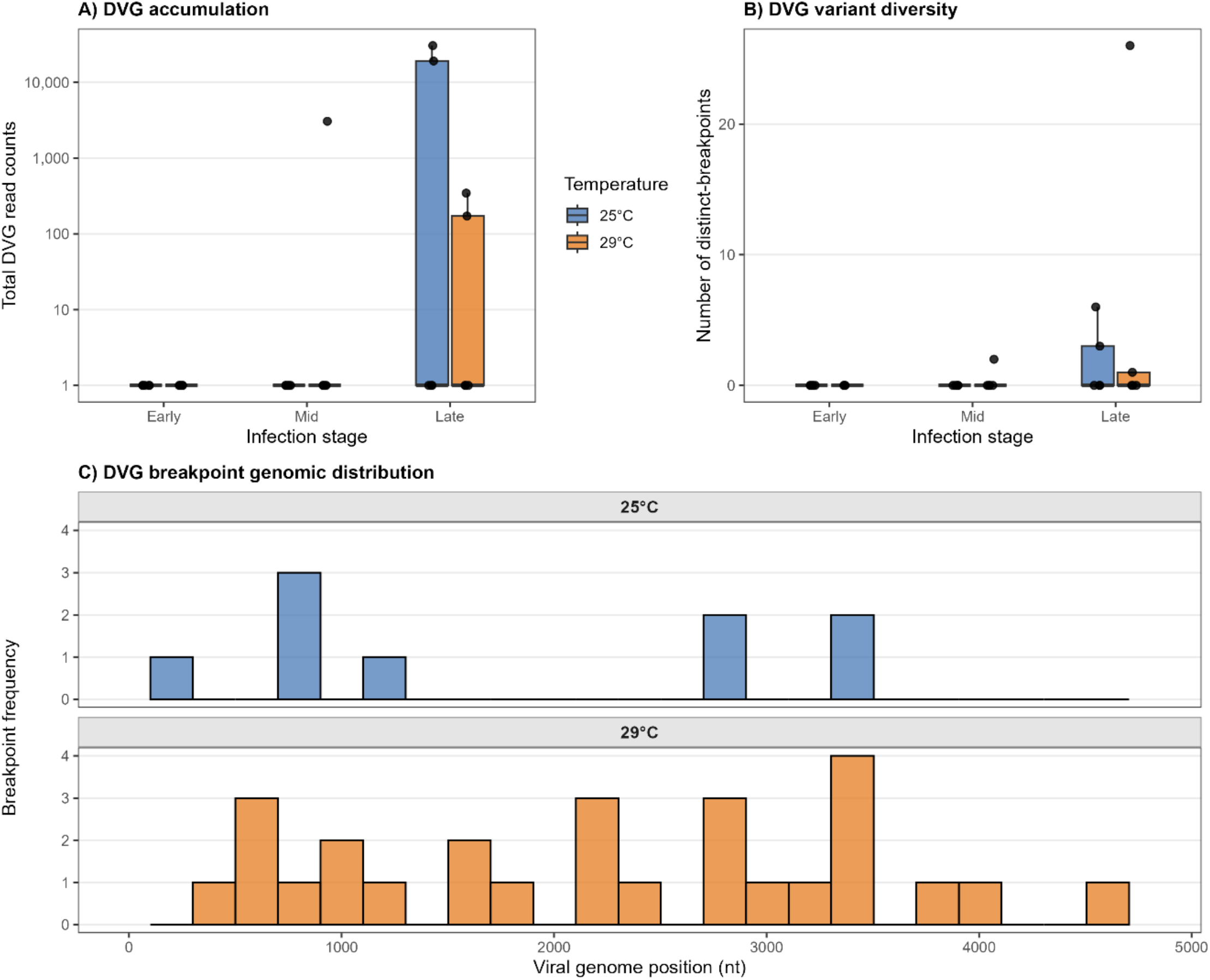
Defective viral genome (DVG) persistence and generation for DAV under standard and thermal stress conditions. **(A)** DVG accumulation quantified as total DVG read counts across infection stages in persistently infected flies maintained at 25°C (blue) and 29°C (orange). **(B)** DVG variant diversity measured as the total number of distinct DVG breakpoint events detected per biological sample (pool of 5 flies) across infection stages. **(C)** Genomic distribution of DVG breakpoint coordinates mapped along the DAV reference genome (4806 nt) at 25°C (top panel, blue) and 29°C (bottom panel, orange).

**Supplementary Figure 4.**
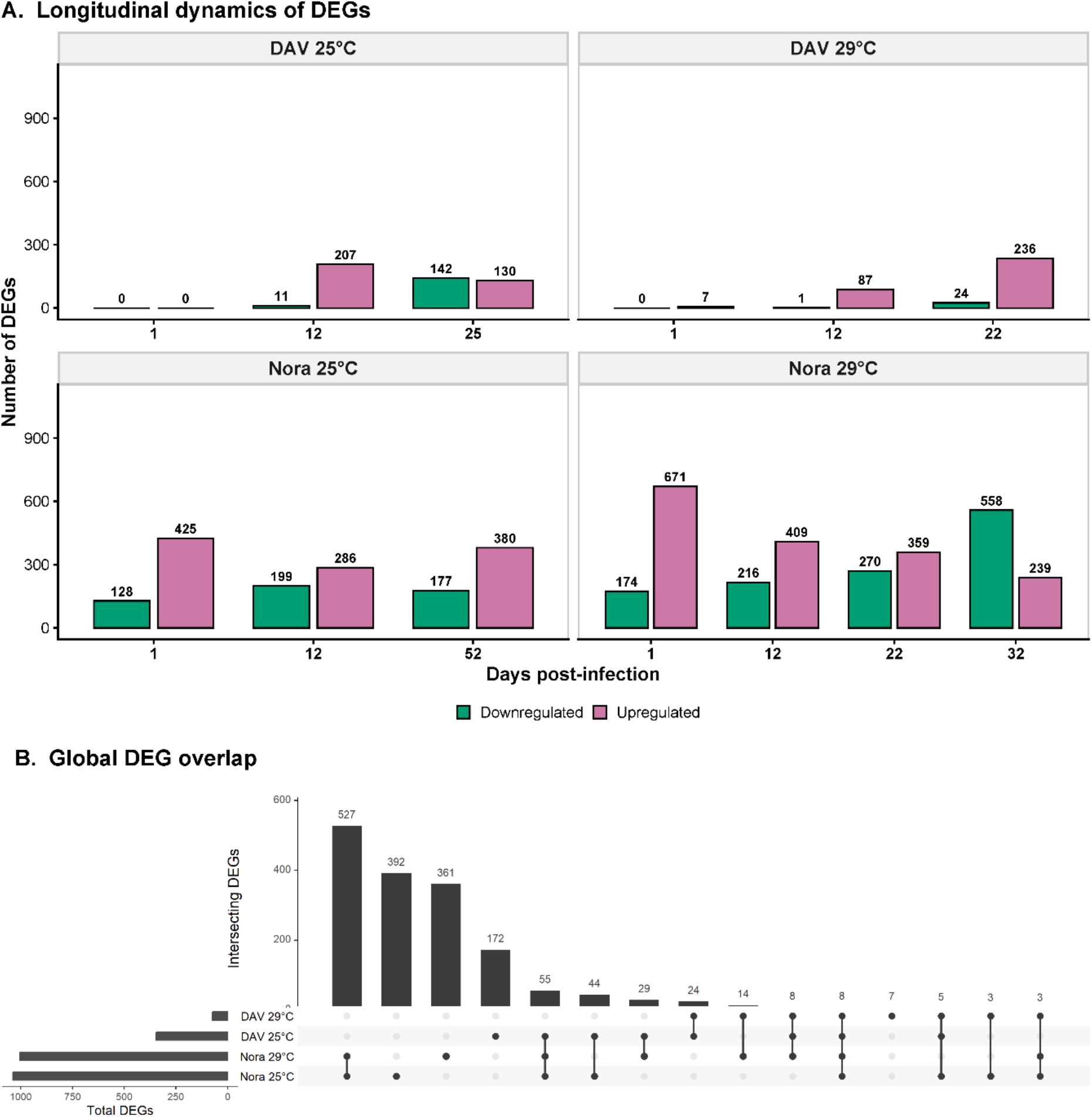
Longitudinal dynamics and global overlap of differentially expressed genes (DEGs) during persistent viral infections across thermal regimes. **(A)** Number of host DEGs (*padj*< 0.05, |log_2_ fold change| > 1.0) across days post-infection, quantified relative to age- and temperature-matched uninfected controls for DAV (top panels) and Nora virus (bottom panels) at 25°C (left panels) and 29°C (right panels). Bars indicate the total number of downregulated (green) and upregulated (purple) genes at each sampled time point, with gene counts displayed above each bar. **(B)** UpSet plot showing the global overlap of cumulative DEGs (pooled across all time points) across viral species and thermal regimes. Vertical bars represent the number of unique or shared DEGs for each combination indicated by the connected black dots in the matrix below.

